# Benchmarking Association Analyses of Continuous Exposures with RNA-seq in Observational Studies

**DOI:** 10.1101/2021.02.12.430989

**Authors:** Tamar Sofer, Nuzulul Kurniansyah, François Aguet, Kristin Ardlie, Peter Durda, Deborah A. Nickerson, Joshua D. Smith, Yongmei Liu, Sina A. Gharib, Susan Redline, Stephen S. Rich, Jerome I. Rotter, Kent D. Taylor

## Abstract

Large datasets of hundreds to thousands of individuals measuring RNA-seq in observational studies are becoming available. Many popular software packages for analysis of RNA-seq data were constructed to study differences in expression signatures in an experimental design with well-defined conditions (exposures). In contrast, observational studies may have varying levels of confounding of the transcript-exposure associations; further, exposure measures may vary from discrete (exposed, yes/no) to continuous (levels of exposure), with non-normal distributions of exposure. We compare popular software for gene expression - `DESeq2, edgeR`, and `limma` - as well as linear regression-based analyses for studying the association of continuous exposures with RNA-seq. We developed a computation pipeline that includes transformation, filtering, and generation of empirical null distribution of association p-values, and we apply the pipeline to compute empirical p-values with multiple testing correction. We employ a resampling approach that allows for assessment of false positive detection across methods, power comparison, and the computation of quantile empirical p-values. The results suggest that linear regression methods are substantially faster with better control of false detections than other methods, even with the resampling method to compute empirical p-values. We provide the proposed pipeline with fast algorithms in R.

## Introduction

Many studies of phenotypes associated with gene expression from RNA-seq consist of small sample sizes (tens of subjects) and are focused on comparisons of transcriptional expression patterns between well-delineated states, such as different experimental conditions, tumor versus non-tumor cells (1; 2), and disease vs non-disease groups (3). Some studies are designed to identify differential expression across hidden, discrete conditions (4). Epidemiological cohorts have recently utilized stored samples to facilitate the use of RNA-seq data in studies of association with subclinical phenotypes such as blood biomarkers, imaging, and other physiological measures, with often continuous measures being used in statistical analyses.

High throughput RNA sequencing enables broad assaying of a sample’s transcriptome (5) and has been in increasing use for over a decade (6). A large variety of analytic and statistical approaches have been developed to address scientific questions such as alternative splicing, differential expression, and more (4; 7–11), often building on methods developed for analyses of expression microarrays (12–14); comprehensive reviews are available (15–19). In this work, we are specifically interested in differential expression analysis with continuous exposures, and we assume that count data are already prepared and available to the analyst. Popular software packages for differential expression analysis include the `DESeq2` R package (9), which models the expression counts as following a negative binomial distribution, with shrinkage imposed on both the mean and the dispersion parameters, based on estimates from the entire transcriptome, or user-supplied values. `EdgeR` (7) uses a negative binomial model similar to the `DESeq2` model for transcript counts, in combination with overdispersion moderation. `EdgeR` was primarily designed for differential expression analysis between two groups when at least one of the groups has replicated measurements (20). `Limma` (21) uses linear models, which are very flexible and can effectively accommodate many study designs and hypotheses. Similar to the `DESeq2` and `edgeR` packages, `Limma` also uses an empirical Bayes method to borrow information across transcripts to estimate a global variance parameter that is applied for the computation of variance parameters of each single transcript. It uses log transformation and weighting, known as the “voom” transformation, in the final linear model that is used for differential expression analysis. We refer to it henceforth as the limma-voom. Prior to differential expression analysis, library normalization is performed (22). Popular approaches are the TMM (trimmed-means of M-values) normalization (23), implemented in `edgeR`, and the size factors normalization (24), implemented in `DESeq2`.

Sleep disordered breathing phenotypes, such as the Apnea-Hypopnea Index (AHI), the number of apnea and hypopnea events per hour of sleep, provides a quantitative assessment of the severity of the disorder, with no clear threshold above which different biological processes occur (although thresholds are used for clinical decision making and health insurance reimbursement). Association analysis with continuous exposures provides different challenges than those traditionally encountered. The distribution of such exposures may have strong effects on the association analysis results, regardless of the underlying associations, due to the combination of skewed exposure distributions and the distribution of RNA-seq read count data, that are generally over-dispersed with occasional extreme values. As observational study data analyses may include covariates, statistical methods from experimental studies (e.g., exact tests) cannot be applied.

In this manuscript, we compare the `DESeq2`, `edgeR`, and `limma-voom` analysis approaches for differential expression analysis, with linear regression–based approaches that do not use the empirical Bayes approach for estimating variance parameter across the transcriptome. We study the computation of p-values using resampling of phenotype residuals, while preserving the structure of the data. This addresses the limitation of permutation noted by others in the context of differential expression analysis of RNA-seq (21), where permutation may not be tuned to test a specific null hypothesis because in its standard form it “breaks” all relationships between the permuted variable and the rest of the dataset. Finally, we study the use of empirical p-values that tune the original p-values based on the residual resampling scheme. Throughout, we use a dataset with sleep disordered breathing phenotypes and RNA-seq from the Multi-Ethnic Study of Atherosclerosis as a case study. We demonstrate the statistical implications of performing association analysis of RNA-seq with continuous, non-normal exposures, compare analysis methods, and develop recommendations.

## Methods

### The Multi-Ethnic Study of Atherosclerosis (MESA)

MESA is a longitudinal cohort study, established in 2000, that prospectively collected risk factors for development of subclinical and clinical cardiovascular disease among participants in six field centers across the United States (Baltimore City and Baltimore County, MD; Chicago, IL; Forsyth County, NC; Los Angeles County, CA; Northern Manhattan and the Bronx, NY; and St. Paul, MN). The cohort has been studied every few years. The present analysis considers N = 462 individuals who participated in a sleep ancillary study performed shortly following the participants Exam 5 during 2010-2013 (25; 26), with RNA-seq measured via the Trans-Omics in Precision Medicine (TOPMed) program. Here, we used RNA-seq data with RNA extracted from whole blood drawn in Exam 5 (2010-2012). Sleep data were collected using standardized full in-home level-2 polysomnography (Compumedics Somte Systems, Abbotsville, Australia, AU0), as described before (26). Of the 462 participants in the current analysis, there were 196 African-Americans (AA), 259 European-Americans (EA) and 125 Hispanic-Europeans (HA). RNA sequencing in MESA is briefly described in the Supplementary Materials.

### Sleep disordered breathing measures

As examples for continuous exposures from population-based studies, we took three sleep disordered breathing measures: (1) the Apnea-Hypopnea Index (AHI), defined as the number of apnea (breathing cessation) and hypopnea (at least 30% reduction of breath volume, accompanied by 3% or higher reduction of oxyhemoglobin saturation compared to the baseline saturation) per 1 hour or sleep; (2) minimum oxyhemoglobin saturation during sleep (MinO2), and (3) average oxyhemoglobin saturation during sleep (AvgO2). We chose these traits because they are clinically relevant, often used in sleep research studies, and represent exposures that may alter gene expression (via hypoxemia and sympathetic activation). The AHI had the least skewed distribution of the considered phenotypes, and AvgO2 had the longest “tail” of small values in the residual distribution. Residuals were obtained by regression the sleep measures on age, sex, body mass index (BMI), study center, and self-reported race/ethnic group.

### Compared tests of associations between exposure and transcripts

We compared the standard packages `DESeq2, edgeR, limma`, and linear regression-based approaches, in which we always applied log transformation on the transcript counts, and then applied linear regression. Because some of the observed transcript count values are zero, which cannot be log transformed, we compared a few approaches for replacing zero values. For a given transcript *j,* denote the minimum observed transcript level that is higher than zero by *m_j_* = min {*t*_*j*1_,…, *t_jn_*: *t_ji_* 0 for *i* – 1,…, *n*}. We compare the following approaches, applied on each transcript *j,j* = 1,…,*k* separately:

A1. SubHalfMin: Replace zero values with 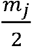
A2.AddHalfMin: Replace all values *t_ij_* by 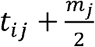
A3.AddHalf: Replace all values *t_ij_* by 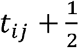.

### Conceptual framework for studying analysis approaches

To study performance of various analysis approaches, we performed simulation studies. Simulation study 1 was used to assess type 1 error across methods when using output p-values, and when using “empirical p-values”, which are p-values that account for true distribution of the p-values under the null and are described later. Simulation study 2 was used to assess power in transcriptome-wide analysis settings, when using methods that control the type 1 error according to simulation study 1. In addition, we performed a simulation study (Supplementary Materials) to assess power for testing of individual transcript according to various distributional characteristics of transcript counts. The goal was to identify approaches for filtering transcripts for association analysis that will optimize power. All simulations used a “residual permutation” (below). The reported criteria for declaring differentially expressed transcripts were False Discovery Rate (FDR) controlling p-values <0.05 based on the Benjamini-Hochberg (BH) procedure, and based on the local FDR procedure implemented in the `qvalue` R package, Family-Wise Error Rate (FWER) controlling p-values <0.05 based on the Holms procedure, and an arbitrary threshold of p-value<10^-5^.

### Residual permutation approach for simulations and for empirical p-value computation

To generate realistic simulation studies in which: (a) the data structure, including the exposure, covariates, and outcome distributions; and (b) their relationships, aside from the exposure-outcome association, are the same as in the real data, we used a residual permutation approach. We regressed each sleep exposure of interest *X* on the covariates ***C*** and estimated their effect ***α***. We then obtained residuals, defined as:

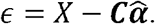

To study type 1 error, we permuted these residuals at random to obtain *perm*(*ϵ*), and generated a sleep exposure unassociated with any of the RNA-seq measures by:

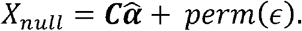

We repeated this procedure 1000 times for evaluating type 1 error control. We generated simulated data under four power simulations in a similar approach, with the difference that we forced a specific correlation value between the simulated sleep exposure and a specific transcript. To this end, for a given transcript *j* measured on individuals *i* = 1,…, *n*, we computed the rank of each individual: *r*(*t*_1*j*_), …, *r_n_*(*t_nj_*). To set a correlation *ρ* between the simulated *X_ρ_* and transcript *j* we sampled *ρ* × *n* (rounded) indices from 1, …,*n*, corresponding to *ρ* × *n* individuals for which we forced their ranks in the permuted residual values, now denoted by *perm*(*ϵ*)_*ρ*_, to be the same as their ranks in the transcript values (note that the transcript values are never changed). For the rest of the individuals, the permuted residuals are completely random. When multiple individuals have the same transcript counts (i.e., their ranks are tied), we randomly assign their ranks. For example, if 100 people have zero counts for a given transcript, each of these individuals will be equally likely to have the rank of 1, 2, …, or 100. The code for generating this residual permutation approach is provided in the Supplementary Information and in a dedicated GitHub repository https://github.com/nkurniansyah/RNA-Seq_continuous_exposure.

### Empirical p-values to account for the null distribution of p-values

We used the residual permutation approach, under the null hypothesis, to generate a null distribution of p-values and to compute empirical p-values. When the distribution of p-values under the null hypothesis is unknown, and specifically when it is not uniform, their values are not reliable for hypothesis testing. Alternative approaches compute “empirical p-values” with the goal of generating an appropriate p-value distribution, i.e., in which an empirical p-value *p_e_* satisfies Pr(*p_e_* < 0.05|*H*_0_) = 0.05 (Supplementary Materials).

For computing empirical p-values, we use a relatively small number of residual permutations (in comparison to the number of permutations used for computing permutation p-values) followed by transcriptome-wide association studies. We use the results of these transcriptome-wide tests under permutation to compute the null distribution of p-values, which is then used to compute the empirical p-values. We compare two types of empirical p-values: quantile empirical p-values, and Storey empirical p-values implemented in the `qvalue` R package (27). The quantile empirical p-value approach is inspired by previously proposed procedures based on permutation (28) of phenotypes (rather than residuals). It estimates the null distribution of p-values non-parametrically, and the quantile empirical p-value is the quantile of the raw p-value in this distribution. The Storey empirical p-values uses the null distribution of the test statistics to identify whether a transcript is likely sampled from the null or a non-null distribution. Both implementations assume that the empirical null distribution is the same for all transcripts. We used 100 residual permutations to compute test statistics and p-values under the null and compared the empirical p-values to standard permutation p-values.

### Resampling approach for binary exposure phenotypes

We compared the analysis of a continuous exposure to that of a dichotomized variable. Instead of a sleep measure, we used body mass index (BMI), because it is known to have large impact of gene expression and is therefore a powerful phenotype for such a comparison. BMI was dichotomized to “obese” if BMI ≥ 30kg/m^2^ and non-obese otherwise. Because obesity is binary and, therefore, the residual permutation approach is not appropriate as proposed for continuous variables, we generated a binomial obesity variable based on BMI probability given covariates. Given a logistic model 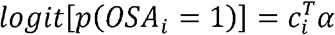, we estimated the covariates’ association parameters 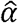 and obtained estimated probabilities for obesity for each person *i* = 1,…, *n* by 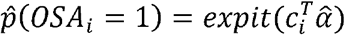. Based on these estimated outcome probabilities, we sampled random obesity status as binomial variables.

## Results

MESA participant characteristics are provided in the Supplementary Material, Table S1. The distribution of the raw phenotypes AHI, MinO2, and AvgO2, and their residuals after regression on covariates is provided in Figure 1, demonstrating the high non-normality. Simulations were performed after normalizing the data so that each library has the same size (prior to filtering), which we set to the median observed value (i.e., median normalization) in the raw reads, or 23,210,672. Results for some of the settings in simulation study 1 under TMM and size factor normalizations are provided in the Supplementary Materials.

**Figure 1:**
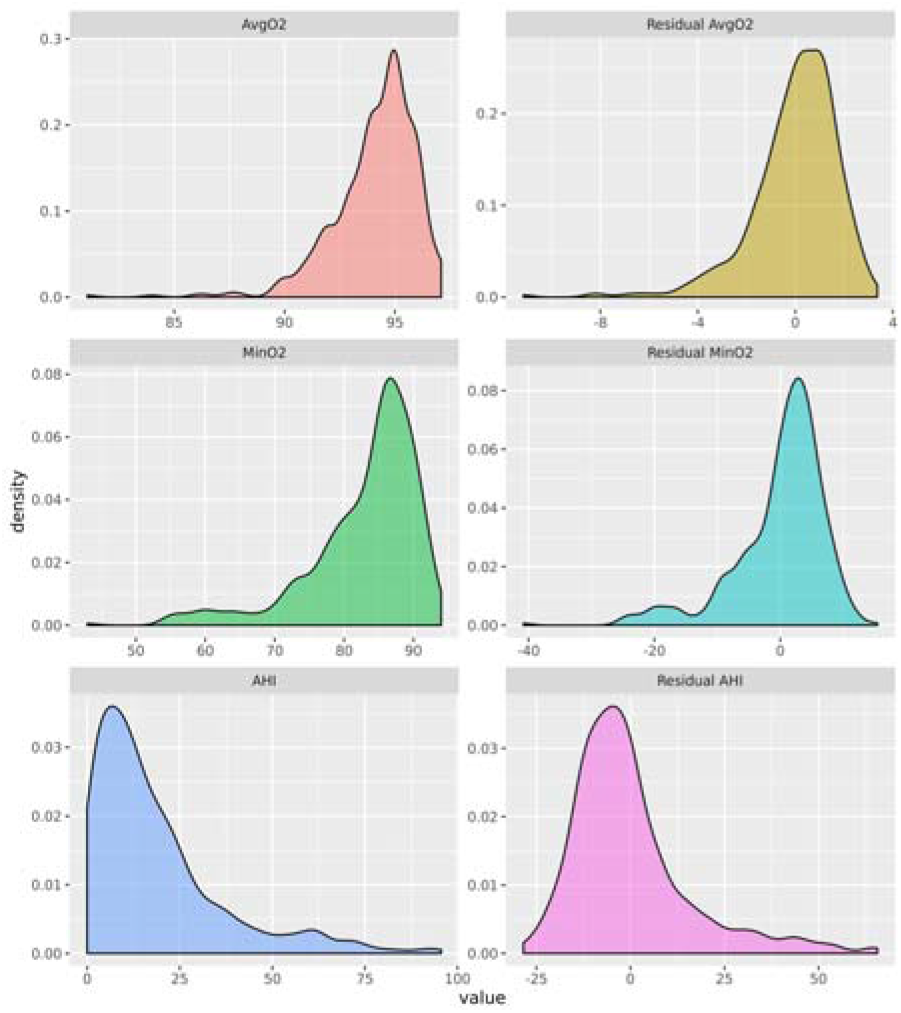
Distributions of the three sleep-disordered breathing exposure phenotypes used as case studies in this manuscript. The left column provides the empirical density functions of the raw phenotypes, the right column provides the empirical density function of their residuals after regressing on age, sex, BMI, self-reported race/ethnic group, and study center. AvgO2: average oxyhemoglobin saturation during sleep. MinO2: minimum oxyhemoglobin saturation during sleep. AHI: Apnea Hypopnea Index.

### Simulation study 1: type 1 error analysis

After normalization, we applied filters to remove lowly expressed transcripts. There were 58,311 transcripts. After applying filters requiring that the (a) maximum read count is ≥10 and that (b) the proportion of individuals with zero counts for a transcript across the sample is not higher than 0.75 (see Supplementary Materials for more information on filters), 23,004 transcripts were available for the simulation study. We used residual permutation to generate simulated SDB phenotypes that are not associated with the transcripts, but maintain the same correlation structure with the transcript and covariates. We generated 100 datasets with simulated SDB phenotypes, and performed analyses. Complete results showing the average number of false positive detection based on the existing packages `limma`, `edgeR`, and `DESeq2`, as well as the three linear regression analyses described here, are provided in Supplementary Figures S3-S5. These results include comparisons of raw p-values, the proposed quantile empirical p-values, and the empirical p-values provided in the `qvalue` R package (27), and for the three SDB phenotypes.

We found that the number of false positives vary with the exposure phenotypes, with analyses of MinO2 (Figure 2) generally resulting in more false positive detections than analyses of the AHI, with intermediate numbers for AvgO2 (Figures S3-S5 in the Supplementary Materials). Figure 2 compares the average number of falsely discovered transcript associations when using simulated sleep phenotypes mimicking MinO2 using the residual permutation approach by focusing on `limma`, `edgeR`, `DESeq2`, and linear regression applied on log2 of expression counts with SubHalfMin. For each method, type I error was determined using raw p-values and Storey empirical p-values, with significance thresholds based on Benjamini-Hochberg (BH) FDR, local FDR (qvalue package), and Holms Family-Wise Error Rate (FWER). Empirical p-values usually reduced the number of false detections, with the method in the `qvalue` package being usually more conservative than the quantile-based empirical p-values method. Compared to linear regression-based approaches, `DESeq2`, `edgeR` and `limma-voom` had many false detections when using the raw p-values, even after applying multiple testing corrections. The three linear regression-based methods described here were quite similar, with the AddHalf approach often resulting in slightly more false detections. Based on these results, we chose to move forward for the next set of simulations with linear regression with SubHalfMin for handling of zero counts.

**Figure 2:**
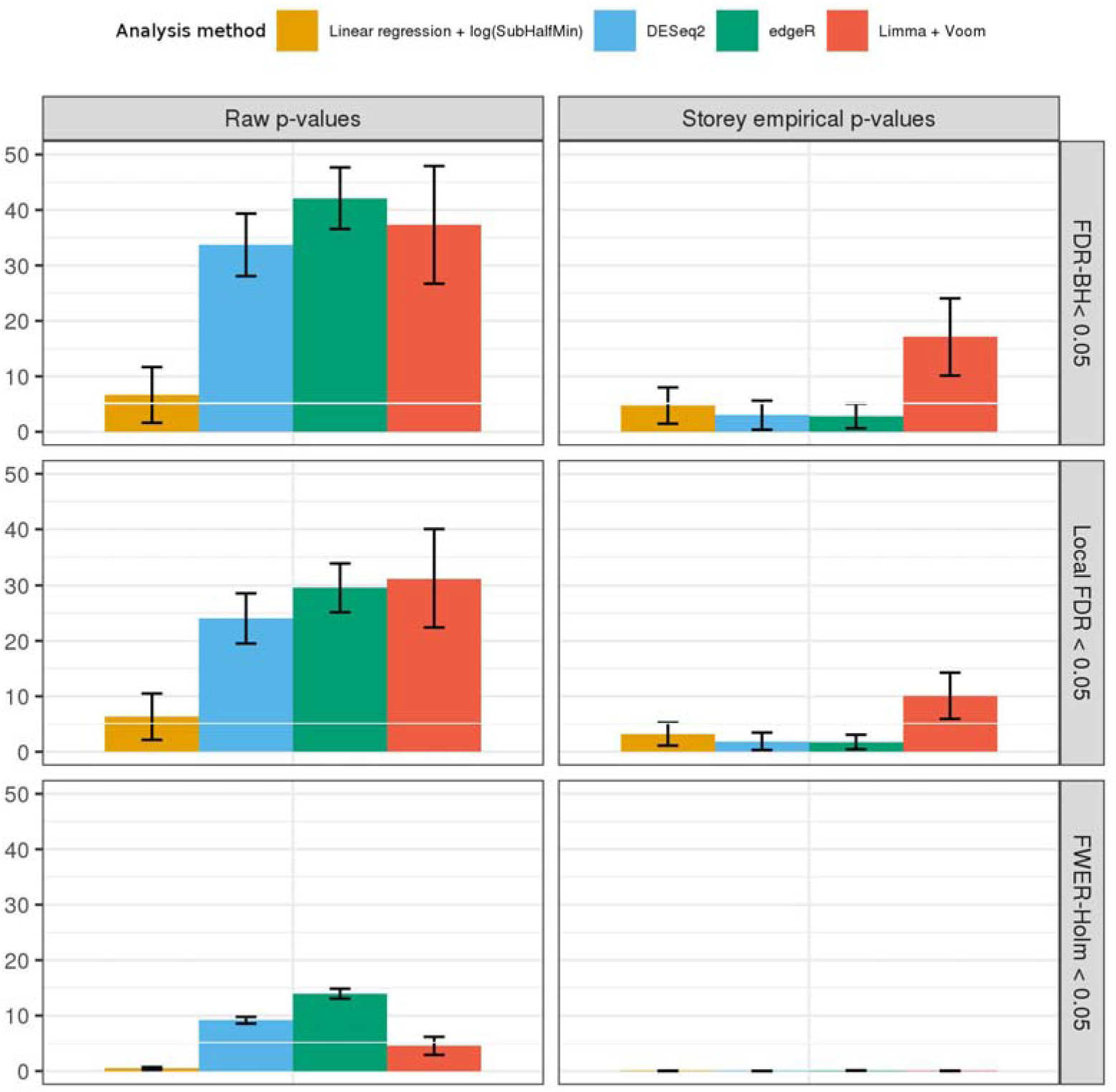
Average number of false positive transcript associations detected by various methods used in simulation study 1 and computed over 100 repetitions. We used the residual permutation approach to mimic the MESA data set with the sleep phenotype MinO2. The methods reported here are linear regression (applied on log2-transformed transcript counts, with zero values replaced with SubHalfMin); `DESeq2 edgeR`, and `limma-voom`. The left column provides results when using raw p-values, the middle corresponds to use of quantile-empirical p-values, and the right corresponds to Storey empirical p-values. We report false positive detections as those with Benjamini-Hochberg (BH) False Discovery Rate adjusted (FDR) adjusted p-value < 0.05, Local FDR <0.05 (`qvalue` package) and with Holms Family-Wise Error Rate (FWER) adjusted p-values < 0.05. Error bars reflect the mean ± standard error. In Supplementary Figures S3-S5, we provide complete results, including for additional sleep phenotypes: AHI and AvgO2.

### Simulation study 2: power analysis

We performed simulations that mimic transcriptome-wide analysis to assess power. Based on simulations comparing power by transcript distributional characteristics (see Supplementary Materials), we only considered 19,742 transcripts for which no more than 50% of the sample had zero counts. We chose two transcripts, and for each of these and each of the sleep phenotypes, we performed 100 simulations in which we used the residual permutation approach to generate association between the sleep phenotype and the transcript with correlation *ρ* = 0.3. We performed transcriptome-wide association analysis using `DESeq2`, `edgeR`, and linear regression with SubHalfMin transformation (`limma-voom` was not used, given its high rate of false positive detections in some of the settings in simulation study 1). For power, we always used empirical p-values (both types) and determined whether the specific transcript of interest passed the significance threshold based on FDR-adjusted (29) empirical p-value < 0.05. Power was defined as the proportion of the simulations in which the associations was significant, and was consistently higher for the linear regression-based approach compared to `DESeq2` or `edgeR`. For linear regression, the quantile empirical p-values performed essentially the same as Storey’s empirical p-values, while Storey’s empirical p-values resulted in substantially higher statistical power when using `DESeq2` and `edgeR`. We illustrate power comparisons in Figure 3 using Storey’s empirical p-values. Power comparisons using quantile empirical p-values are provided in the Supplementary Materials Figure S8.

**Figure 3:**
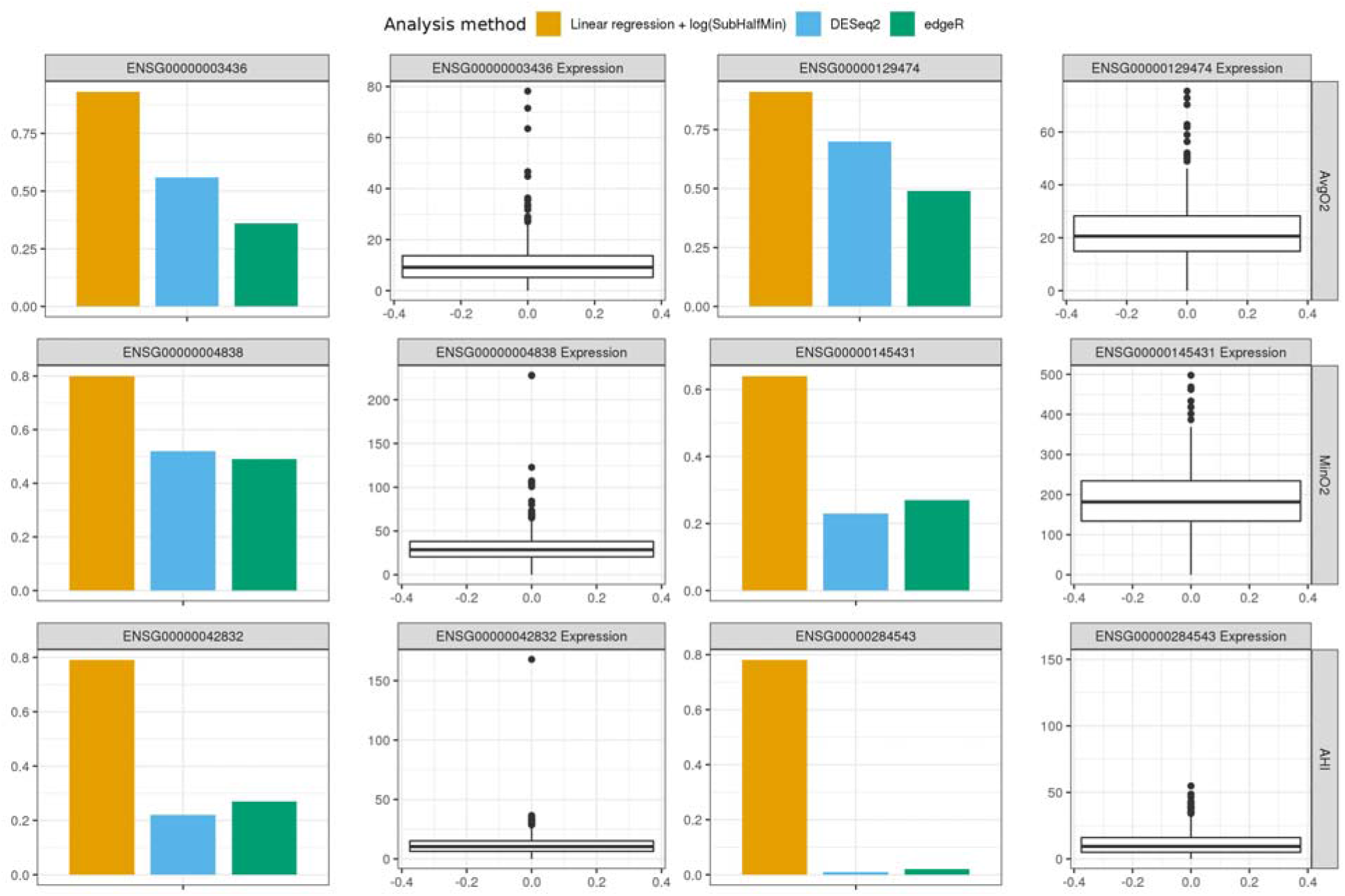
Estimated power for detecting a transcript simulated as associated with the three sleep traits when using Storey empirical p-values, and association is determined significant if its BH FDR-adjusted p-value is <0.05. The transcripts were randomly selected out of available transcripts (after filtering of transcripts with 50% or higher zero counts across the sample). We compared linear regression, `DESeq2`, and `edgeR` in transcriptome-wide association analysis for each of the sleep phenotypes. For each transcript used in simulations, we show both power and the box plot of its distribution in the sample after Median normalization.

### Proposed analysis approach

Based on the above simulation studies, we developed an analytic pipeline as depicted in Figure 4: (a) the raw read count are normalized; (b) filters are applied to remove lowly expressed transcripts and those for which the statistical power is low, as determined by simulations, (c) AddHalfMin transformation is applied for each individual separately, then log transformation is applied on all transcripts, (d) association analyses is performed using linear regression to compute effect sizes and p-values, (e) permutations are computed 100 times on exposure residuals after regressing on covariates, to generate simulated traits that maintain the data structure, (f) each of 100 vectors of simulated traits are analyzed using the same approach as the raw trait, generating p-values, (g) p-values from the analysis of the 100 simulated traits are combined to generate an empirical null distribution of p-values, that are used to generate empirical p-values for the raw trait using the `qvalue` package, and (h) multiple testing correction is applied on the empirical p-values.

**Figure 4:**
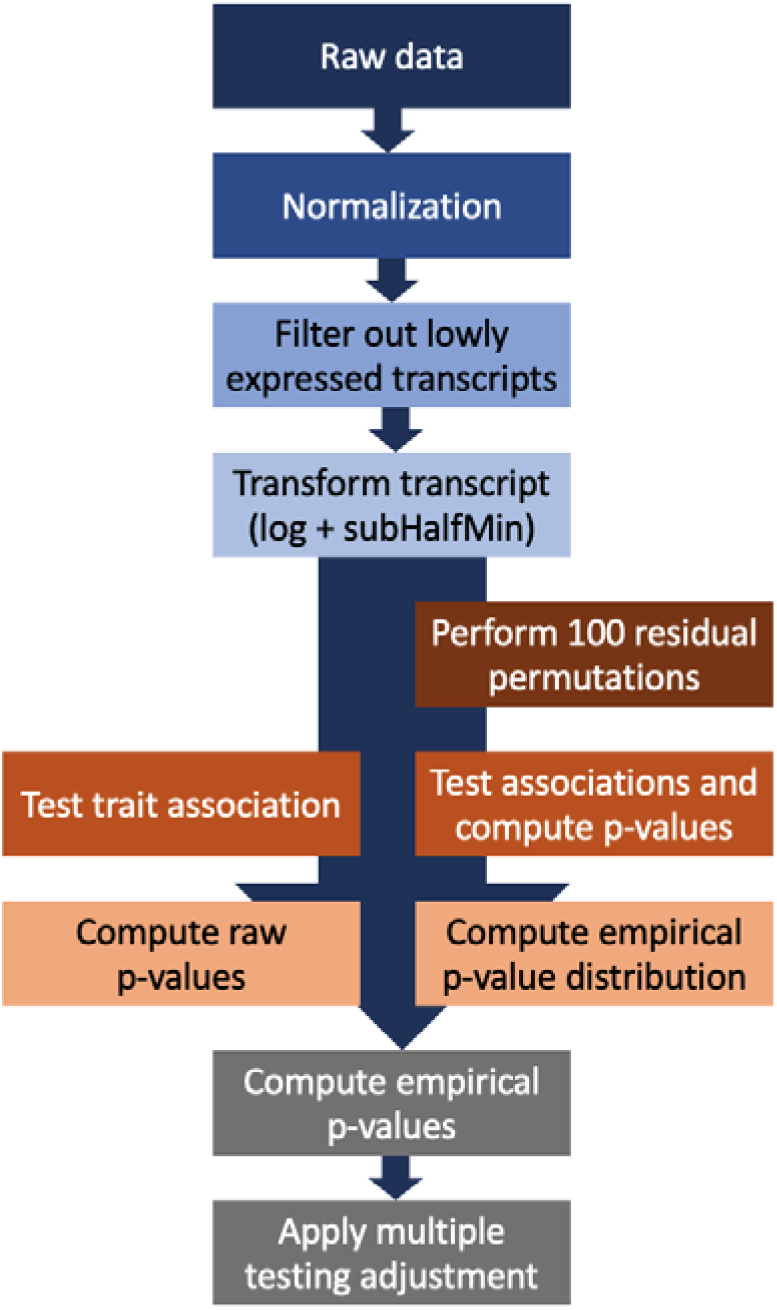
Analysis pipeline for association transcriptome-wide association analysis of continuous exposure phenotypes. The raw data is normalized using library-size normalization, followed by filtering of transcripts, transformation of transcript expression values, then single-transcript testing to obtain raw p-values. In parallel, residual permutation is applied under the null 100 times, and p-values are used to construct an empirical p-value distribution under the null, and to compute empirical p-values. Finally, the quantile empirical p-values are corrected for multiple testing.

### Comparison of analysis of continuous BMI with analysis of dichotomous obesity status

We compared the differential expression of transcripts in analysis of BMI and obesity. Residual permutation procedure was used and quantile-empirical p-values generated for both analyses. A total of 925 MESA individuals had BMI measure available and, for analysis, at least 50% non-zero transcripts were required. For obesity, several non-zero transcript thresholds were examined: 50%, 40%, and 30%. The results were similar for all thresholds, resulting in many more identified transcript associations (446 vs. 251) with continuous BMI compared to using a dichotomous trait (Supplementary Information Figure S9).

### Computing time comparison

The compute time for transcriptome-wide association study was obtained for analyses using `DESeq2`, `edgeR`, and our linear regression implementation. Using our linear regression implementation on a single core, a single transcriptome-wide association study applied on ~19K transcripts and N=462 individuals took less than a minute; when 100 transcriptome-wide association studies applied to residual permutations were included to compute empirical p-values, the time reached 7 minutes, and the maximum memory used was 1.3GB. In comparison, `DESeq2` took 53.5 minutes and `edgeR` took 18.8 minutes for a single transcriptome-wide association study. The maximum memory used for `DESeq2` and `edgeR` was similar at 3.1GB.

### R package

Code for implementing the proposed procedure and for a shiny app is provided in the GitHub repository https://github.com/nkurniansyah/Olivia. The code also provides test of multiple exposure variable at the same time, which applies the multivariate-Wald test, and an efficient implementation of a permutation test when considering a single transcript, rather than a transcriptome-wide analysis. The repository also includes code used for simulations.

### Data availability

MESA data are available through application to dbGaP. Phenotypes are available in MESA study accession phs000209.v13.p3, and RNA-seq data has been deposited and will become available through the TOPMed-MESA study accession phs001416.v2.p1.

## Discussion

We systematically assessed the approaches for studying the association of gene expression, estimated using RNA sequencing, with continuous and non-normally distributed exposure phenotypes. We found that linear regression-based analysis performs well for continuous phenotype associations, and is computationally highly efficient. We used a residual permutation approach to study the distribution of p-values under the null of no association between the phenotypes and RNA-seq, and used this approach to further study power, and to compute empirical p-values. Notably, the residual permutation approach allows for the dataset to have the same correlation structures and associations between the phenotypes and the transcripts and covariates, while eliminating the transcript-phenotype associations. We implemented this approach in an R package and developed an R shiny app, to make our pipeline easily accessible to the research community.

Recently, van Rooij et al (30) also performed a benchmarking study comparing analysis approaches for transcriptome-wide analysis of RNA-seq in population-based studies, including when using continuous phenotypes in association testing. While we used similar statistical methods to theirs, we took a different analytical approach. Van Rooij et al. used multiple datasets to apply association analysis between a phenotype and transcripts, and assessed replication between analyses. We, on the other hand, leveraged simulations to generate data under a known association structure. In addition, we were motivated by a specific problem: highly non-normal sleep exposure measures, often leading to suboptimal control of Type 1 error. Thus, it was critical to assess control of false discovery under the null hypothesis. Notably, sleep phenotypes are less often available and there are no other large observational studies data sets to our knowledge with both RNA-seq measures and similar SDB phenotypes. Some of our findings are similar to those of van Rooij et al.: they also recommend using linear regression analysis, and they also found that using a continuous phenotype is generally more powerful than dichotomizing it (in agreement with what is known from statistical literature). Similarly, they found that normalization method had very little effect on the results. However, they recommend testing all genes, while we recommend filtering transcripts with at least 50% zero counts, based on our power simulations. Additional future work is needed to evaluate various filtering criteria, and to develop methods that allow for flexible, non-linear modeling of the association between phenotype and gene expression while remaining computationally efficient to allow for permutation analysis.

We propose to compute p-values under the null hypothesis of no association between the transcript and the exposure phenotype by permuting residuals of the exposure phenotype after regressing on covariates, and re-structuring the exposure by summing the permuted residuals with the estimated mean, and thus maintain the overall data structure except for the exposure-outcome association of interest. Outside the gene expression literature, others have proposed to permute residuals rather than the outcome. For example, previous permutation methods proposed to permute residuals of the outcome after regressing on covariates (31), or to permute the residuals of the exposure phenotypes without constructing a new exposure phenotype by summing the permuted residuals with the estimated mean (32). It will be interesting to perform a more comprehensive study of statistical permutation approaches for RNA-seq association analyses, as well as studying them in the context of mixed models.

We recommend using empirical p-values, which require 100 residual permutation, and therefore, performing 101 transcriptome-wide association analyses instead of one. Considering Figures S3-S5 in the Supplementary Information, one can see that in most settings, linear regression methods do not have many false positive detections even when raw p-values are used. However, we chose to be more conservative by strongly protecting the analysis from false positive detections. Importantly, the linear regression analysis with empirical p-values had higher power than the other common approaches (`DESeq2, edgeR`), indicating simultaneous improvement in controlling false positives and increasing power. Unfortunately, we cannot effectively estimate the FDR in these simulations. FDR is defined as the proportion of false discoveries out of all discovered (significant) associations. In simulation study 1, none of the transcripts were associated with the outcomes, so that any estimated FDR would be 100%. Under the alternative, one can suggest to use the number of wrongly discovered associations to estimate the FDR.

However, many transcripts are highly correlated with the one simulated to be associated with the exposures, and are therefore associated with the exposure by design, and thus the number of transcripts falsely detected as associated with the exposure cannot be easily determined.

The empirical p-values procedure uses p-values from the entire tested transcriptome to compute the empirical null distribution. This encapsulates the assumption that the null distribution of p-values is the same for all transcripts, which is generally a limitation, but has been shown to be often acceptable since it will lead to less power, rather than increasing the number of false detections (33; 34). An approach that does not require this assumption estimates the null distribution for p-value for each transcript separately, which is a standard permutation approach. We investigated this issue by comparing the quantile empirical p-values with the permutation p-values that use 100,000 residual permutations to estimate the null distribution of the p-value of each transcript separately (Figure S2 in the Supplementary Materials). The two p-value distributions are very similar. Therefore, a computationally expensive permutation approach, as well as other approaches proposed by investigators, such as estimating null distributions across sets of transcripts with similar properties (33; 35), are likely unnecessary and not superior to the computationally efficient empirical p-values method. Another approach for estimating the null distribution of p-values uses the primary results, without any permutation (36; 37). These approaches also use the assumption that the null p-value distribution is the same across transcripts (i.e. a shared null distribution exists). Given the computationally fast implementation of the transcriptome-wide association study, we believe that using residual permutation is beneficial because it allows for a more precise quantification of the null p-value distribution.

Batch effects are important to account for in studies of RNA-seq. Here, we did not study their effect because it was beyond the scope of our investigation. van Rooij et al (30) in their benchmarking study focusing on replication across cohorts, compared a few approaches for adjusting for technical covariates, including estimating and adjusting for latent confounders (38). They concluded that inclusion of more technical adjusting covariates, including hidden confounders, increases the rate of replication between studies.

To summarize, we highlighted the problem of high false positive findings in RNA-seq data when studying the association of continuous exposure phenotypes that are highly non-normal. We developed a computationally efficient pipeline to address the false positive detection problem, and studied strategies to optimize statistical power. Our approach will be particularly useful for epidemiological studies with RNA-seq data that were not designed as disease-focused case-control studies.

## Acknowledgements

This work was supported by the National Heart Lung and Blood Institute grant R35HL135818. MESA and the MESA SHARe projects are conducted and supported by the National Heart, Lung, and Blood Institute (NHLBI) in collaboration with MESA investigators. Support for MESA is provided by contracts 75N92020D00001, HHSN268201500003I, N01-HC-95159, 75N92020D00005, N01-HC-95160, 75N92020D00002, N01-HC-95161, 75N92020D00003, N01-HC-95162, 75N92020D00006, N01-HC-95163, 75N92020D00004, N01-HC-95164, 75N92020D00007, N01-HC-95165, N01-HC-95166, N01-HC-95167, N01-HC-95168, N01-HC-95169, UL1-TR-000040, UL1-TR-001079, UL1-TR-001420. Also supported in part by the National Center for Advancing Translational Sciences, CTSI grant UL1TR001881, and the National Institute of Diabetes and Digestive and Kidney Disease Diabetes Research Center (DRC) grant DK063491 to the Southern California Diabetes Endocrinology Research Center.

Molecular data for the Trans-Omics in Precision Medicine (TOPMed) program was supported by the National Heart, Lung and Blood Institute (NHLBI). RNA-Seq for “NHLBI TOPMed: Multi-Ethnic Study of Atherosclerosis (MESA)” (phs001416.v1.p1) was performed at the Northwest Genomics Center (HHSN268201600032I). Core support including centralized genomic read mapping and genotype calling, along with variant quality metrics and filtering were provided by the TOPMed Informatics Research Center (3R01HL-117626-02S1; contract HHSN268201800002I). Core support including phenotype harmonization, data management, sample-identity QC, and general program coordination were provided by the TOPMed Data Coordinating Center (R01HL-120393; U01HL-120393; contract HHSN268201800001I). We gratefully acknowledge the studies and participants who provided biological samples and data for TOPMed.

## Author contributions

TS conceptualized and drafted the manuscript and supervised the analysis. NZ performed all statistical analysis and data visualization and developed the R package and R shiny app. DN, and JS performed RNA sequencing. FA and KA generated the MESA processed the sequenced RNA to generate the RNA-seq dataset. PD, YL, SSR, JIR, and KDT designed the RNA-seq study in MESA. SR designed and supervised the MESA sleep ancillary study. All authors critically reviewed and approved the manuscript.

