## Supplementary materials for "Benchmarking Association Analyses of Continuous Exposures with RNA-seq in Observational Studies"

Supplementary Information

Tamar Sofer, Nuzulul Kurniansyah, François Aguet, Kristin Ardlie, Peter Durda, Deborah A. Nickerson, Joshua D. Smith, Yongmei Liu, Sina A. Gharib, Susan Redline, Stephen S. Rich,
Jerome I. Rotter, Kent D. Taylor

### RNA sequencing in MESA

RNA-seq was generated from peripheral blood mononuclear cells (PBMCs) of MESA participants obtained in the fifth clinic visit. PBMC samples were sequenced at the Broad Institute (n=498), and at the North West Genomics Center (NWGC; n=468). Both centers used harmonized protocols. RNA samples quality was assessed using RNA Integrity Number (RIN, Agilent Bioanalyzer) prior to shipment to sequencing centers. QC was re-performed at sequencing centers by RIN analysis at the NWGC and by RNA Quality Score analysis (RQS, Caliper) at the Broad Institute. A minimum of 250ng RNA sample was required as input for library construction, performed using the Illumina TruSeq^TM^ Stranded mRNA Sample Preparation Kit. RNA was sequenced as 2x101bp paired-end reads on the Illumina HiSeq 4000 according to the manufacturer’s protocols. Target coverage was of ≥40M reads. Comprehensive information about the RNA-seq pipeline used for TOPMed can be found in <https://github.com/broadinstitute/gtex-pipeline/blob/master/TOPMed_RNAseq_pipeline.md> under MESA RNA-seq pilot commit 725a2bc. Here we used transcript-level expected counts quantified using RSEM v1.3.0 (1).

### Characteristics of MESA participants

Table S1: Characteristics of participants from MESA with RNA-seq and sleep study, by sex.

| Characteristic | Females | Males |
| --- | --- | --- |
| n | 250 | 212 |
| Race (%):  White  Black  Hispanic | 108 (43.2)  57 (22.8)  85 (34.0) | 92 (43.4)  41 (19.3)  79 (37.3) |
| Age (mean(SD)) | 68.37 (9.44) | 67.56 (9.30) |
| BMI (mean (SD)) | 30.73 (5.90) | 28.79 (4.57) |
| AvgO2 (mean (SD)) | 94.24 (1.78) | 94.00 (2.05) |
| AHI (mean (SD)) | 15.41 (15.22) | 22.36 (20.10) |
| MinO2 (mean (SD)) | 82.87 (8.39) | 83.04 (7.56) |

### Study of filtering based on distributional characteristics of transcripts

We studied multiple approaches to filter transcripts and reduce the number of transcripts in the association analysis. The goal was to identify distributional characteristics of the transcripts that result in low power. Based on these, one may be able to retain a smaller number of genes in the analysis and increase power by reducing the multiple testing burden. We note that filtering also affects type 1 error, because the overall distribution of p-values under the null changes depending on the set of transcripts used.

1. **Considered filters**

The considered filters (and characteristics) were, as conditions applied on transcript $j$:

F1. Expression sum filter: The sum of expression counts across all people need to satisfy $\sum_{i=1}^{n} t_{ij}>C_{1}$.
F2. Median filter: The median expression count across all people need to satisfy $median\left( t_{1j},\ldots, t_{nj} \right)>C_{2}$.
F3. Max to Median Ratio filter: The maximum to median expression count across all people need to satisfy: $\left[ \frac{median\left( t_{ij}, i=1, \ldots n \right)}{\max\left( t_{ij}, i=1, \ldots n \right)} \right] >C_{3}$.
F4. Maximum filter: The maximum expression count across all people need to satisfy: $max\left( t_{1j},\ldots, t_{nj} \right)>C_{4}$.
F5. Range filter: The range of read counts for a transcript needs to satisfy:
    $\left( t_{max,j}-t_{min,j} \right)>C_{5}$, where $t_{max,j}$and $t_{min,j}$ are the highest and lowest read counts observed for this transcript across all people in the sample.
F6. Proportion zero count filter: The proportion of individuals with zero expression count needs to satisfy $\frac{1}{n}\sum_{i=1}^{n} 1(t_{ij}=0)\leq C_{6}.$
F7. Coefficient of variation filter: The standard deviation divided by the mean expression value $\frac{sd\left( t_{1j},\ldots, t_{nj} \right)}{mean\left( t_{1}, \ldots, t_{nj} \right)}=C_{7}$need to be within a specified range.
2. **Simulations of transcript characteristics and power**

For each transcript, we used the exposure phenotype AvgO2 to generate residuals, followed by the residual permutation scheme with correlation parameter $\rho=0.3$ to generate a simulated exposure variable that is associated with the transcript. We tested the phenotype-transcript association using linear regression after log applied on SubHalfMin transformation. We repeated the simulations 100,000 times, and computed power for each transcript based on the proportion of simulations in which the raw p-value from testing the association of the transcript with the simulated phenotype was smaller than ${10}^{-6}$. Supplemental File 1 provides results from this power assessment. Based on these simulations, only one clear pattern emerges: a high number of zero expression values across samples for a given transcript leads to low power. Based on this, we proceeded requiring that at least 50% of samples have non-zero values for a transcript for it to be included in the analysis, or equivalently, median expression values higher than 0. Figure S1 demonstrates the loss in power for analyzing a continuous phenotype when the median expression value is 0.

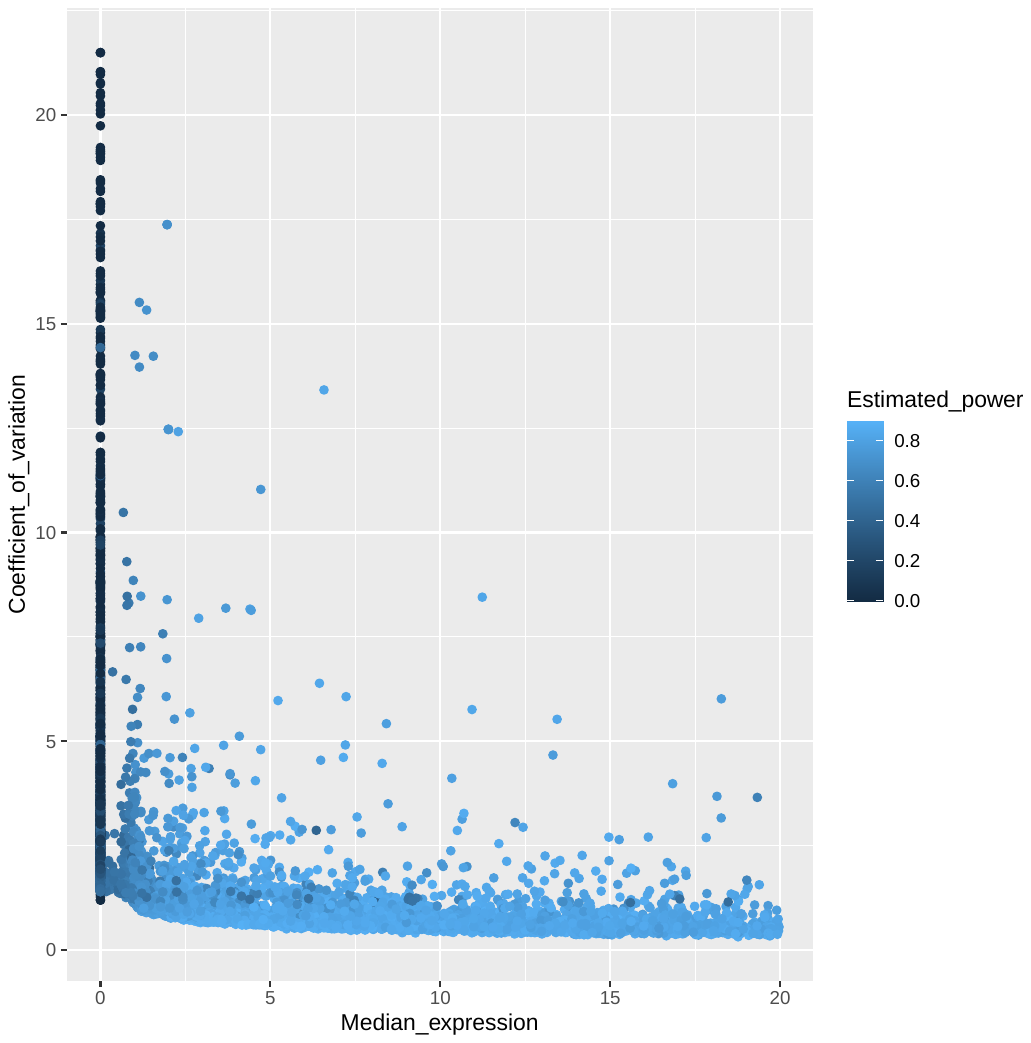

Figure S1: Estimated power to detect associations using linear regression analysis for a given transcript as a function of median expression value (after normalization) and coefficient of variation. Median expression values were capped at 20 in the figure. Power was estimated in 1,000 simulations using the residual permutation approach with $\rho=0.3$, and 100,000 residual permutations with no associations were used to compute permutation p-values for each transcript.

### Permutation and empirical P-values

A standard, well-known approach to computing p-values when the null distribution cannot be specified is permutation. We refer to these p-values as “permutation p-values” and here we provide details in order to contrast them with empirical p-values. We note here that in the statistical literature permutation p-values are sometimes called empirical p-values, but here we use the term empirical p-value as often applied in the gene expression analysis literature.

1. **Permutation p-values**

The idea is that for a given transcript $j$, the transcript expression values, or the exposure (or the residuals) are permuted $B$ times across individuals, and a p-value is computed from each study permutation $p_{b}^{j}, b=1, \ldots, B.$ Then, the permutation p-value is

$$p_{perm}^{j}= \frac{1}{B}\sum_{b=1}^{B} 1\left( p_{b}^{j}<p_{true}^{j} \right),$$

assuming that each $p_{b}^{j}, b=1, \ldots, B$ is drown from the null distribution of $p_{true}^{j}$. The challenge with permutation p-values for transcriptomics is the computation burden. The number of permutations $B$ has to be very large, because the permutation p-values $p_{b}^{j}$ are based on evaluations on the specific transcript permutation. In other words, the entire transcriptomics analysis has to be performed $B$ times, where $B$ is usually at least 100,000, leading to a huge computational burden.

2. **Empirical p-values**

We study the use of *empirical p-values,* computed while capitalizing over all transcripts for computation of the empirical p-value of every specific transcript. We use a non-parametric, quantile-based approach (2), and the method implemented in the qvalue R package (3), both using the residual permutation approach to obtain the empirical distribution of p-values under the null. The specific implementation of the quantile empirical p-values is as follows: Consider the distribution of permutation p-values for a transcript $j,$ defined by $\{ p_{1}^{j}, \ldots, p_{B}^{j}\}$, as $F_{j}\left( x \right)=\frac{1}{B}\sum_{b=1}^{B} 1\left( p_{b}^{j}<x \right).$ Considering all transcripts $j=1, \ldots, k$ passing the filtering criteria and participating in the analysis. Suppose we permute each one of them $B$times, and estimate an empirical p-value distribution as

$$F_{emp}\left( x \right)= \frac{1}{B\times k}\sum_{j=1}^{k} \sum_{b=1}^{B} {1(p}_{b}^{j}<x).$$

Then, if $F_{j}\left( x \right)\approx F_{emp}(x)$, we can use $F_{emp}$ rather than $F_{j}$ to compute empirical p-values rather than permutation p-values, with a substantially smaller $B$, meaning with many less transcriptome-wide association analyses.

3. **Comparison of permutation p-values and empirical p-values**

We compared the quantile empirical p-values to permutation p-values using the three sleep exposures. We removed all transcripts with maximum counts of 10 and more than 50% zero counts in the sampled, and applied median normalization. We then used the residual permutation approach to generate data for simulation under the null of no phenotype-transcript association, and tested for differential using linear regression after log transformation of subHalfMin approach for handling zero counts. We performed permutation analysis $B=100,000$ times ($B$transcriptome-wide residual permutation analyses), and compared the resulting permutation p-values to the empirical p-values obtained using $B=100$ transcriptome-wide residual permutation analyses. Figure 2 provide the comparison, demonstrating that the two p-values are very similar, therefore, it is appropriate to use empirical p-values which are computationally much faster to compute.

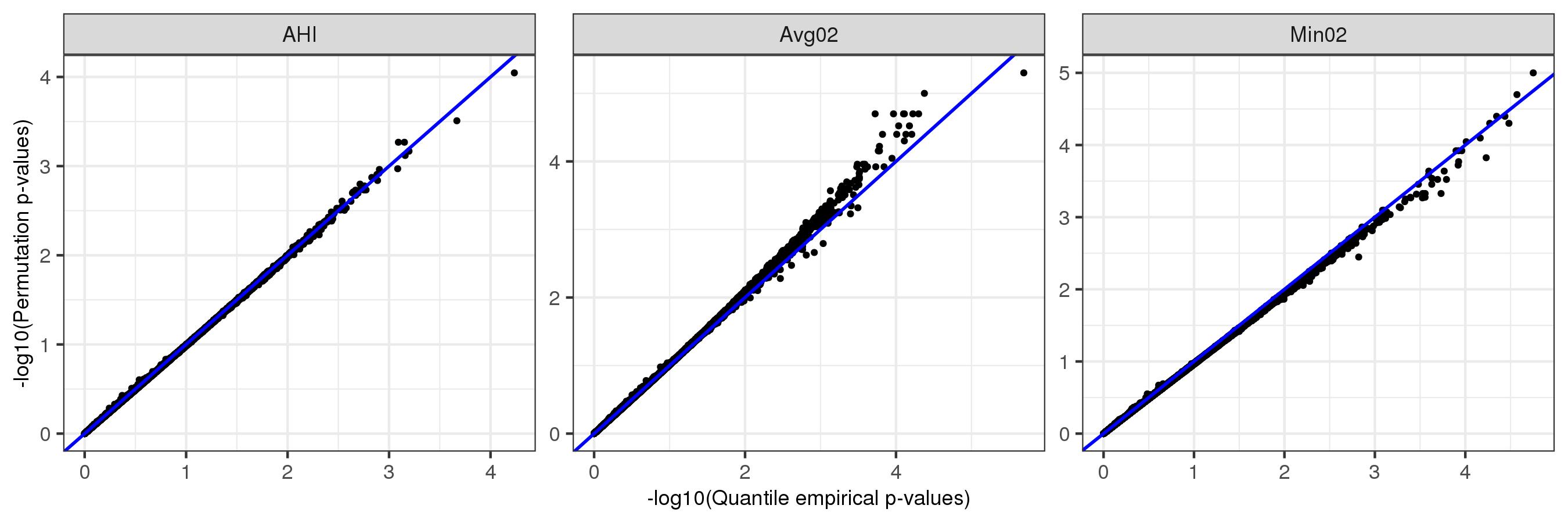

Figure S2: Quantile empirical p-values (computed using $B=100$ residual permutations) versus standard permutation p-values (computed using $B=100,000$ residual permutations) in simulations.

### Results from simulation study 1: assessment of false positive detections

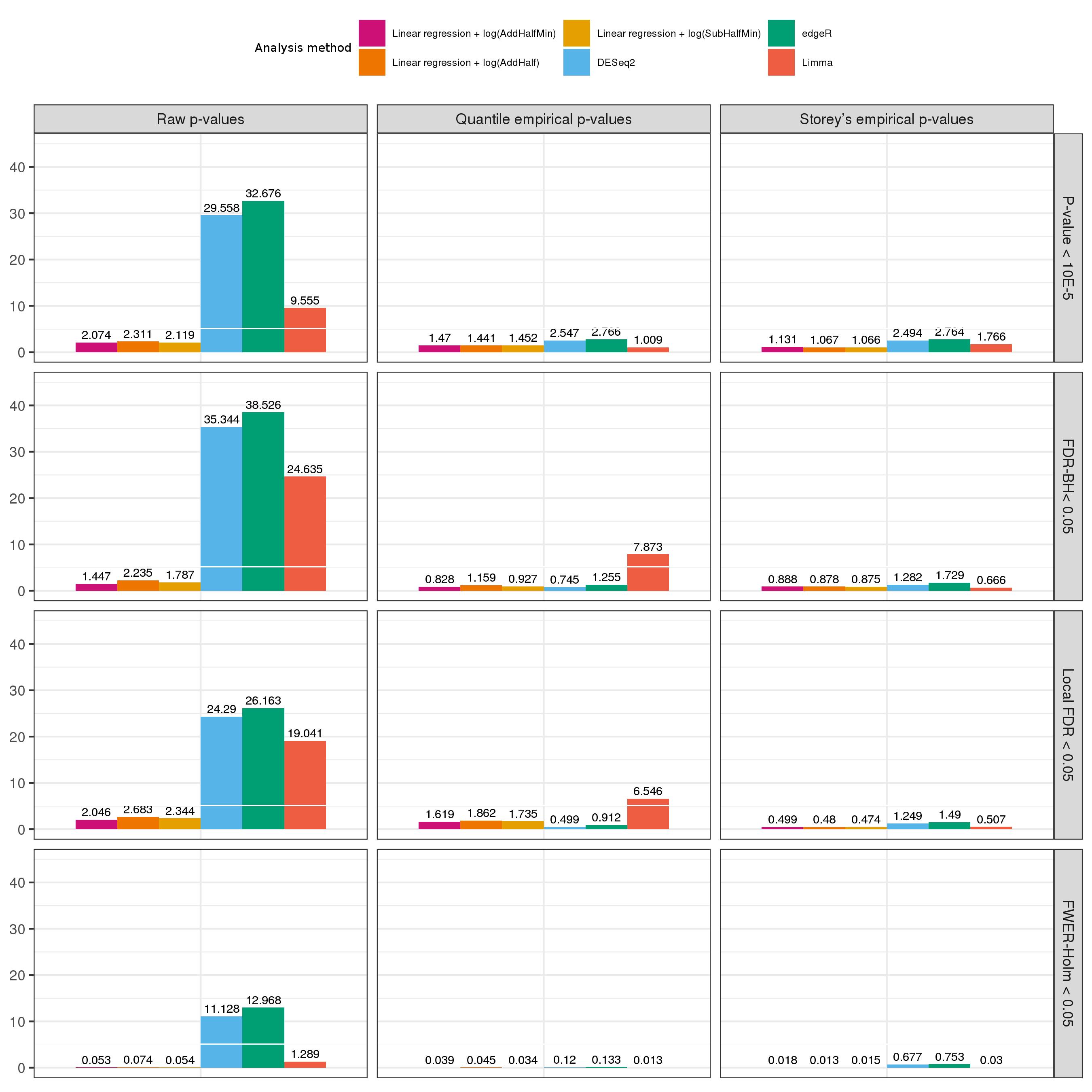

Figure S3: Average number of falsely-detected transcript association with the residual-permuted **AHI** phenotype, estimated over 100 permutations. We compared approaches based on linear regression, DESeq2, limma, and EdgeR packages; raw p-values, quantile-empirical p-values, and Storey’s empirical p-values; and associations declared as significant according the arbitrary threshold of p-value < ${10}^{-5}$, FDR adjustment using the Benjamini-Hochberg (BH) procedure and using the local FDR procedure implemented in the qvalue R package, and FWER adjustment using Holm procedure.

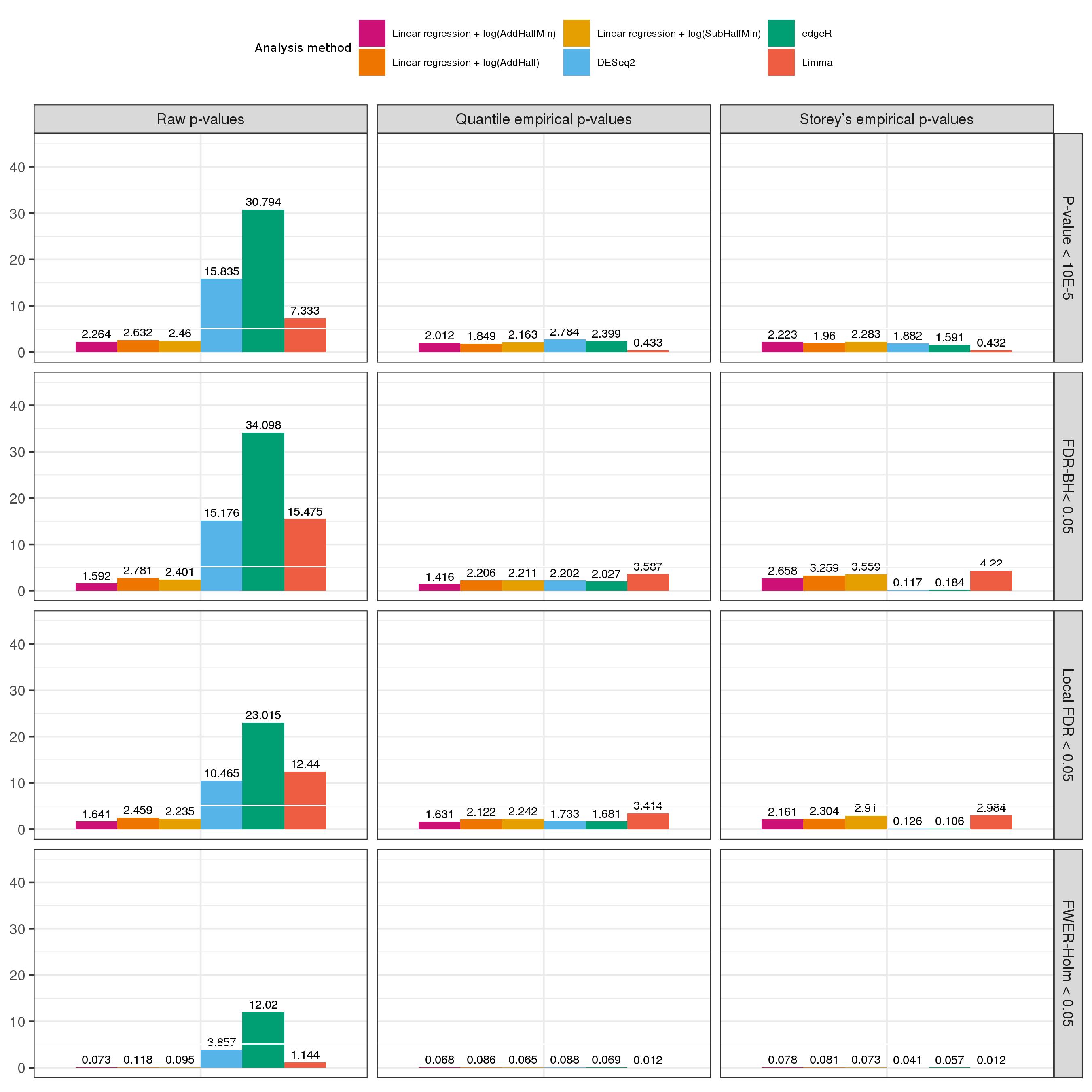

Figure S4: Average number of falsely-detected transcript association with the residual-permuted **AvgO2** phenotype, estimated over 100 permutation. We compared approaches based on linear regression, DESeq2, limma, and EdgeR packages; raw p-values, quantile-empirical p-values, and Storey’s empirical p-values; and associations declared as significant according the arbitrary threshold of p-value < ${10}^{-5}$, FDR adjustment using the Benjamini-Hochberg (BH) procedure and using the local FDR procedure implemented in the qvalue R package, and FWER adjustment using Holm procedure.

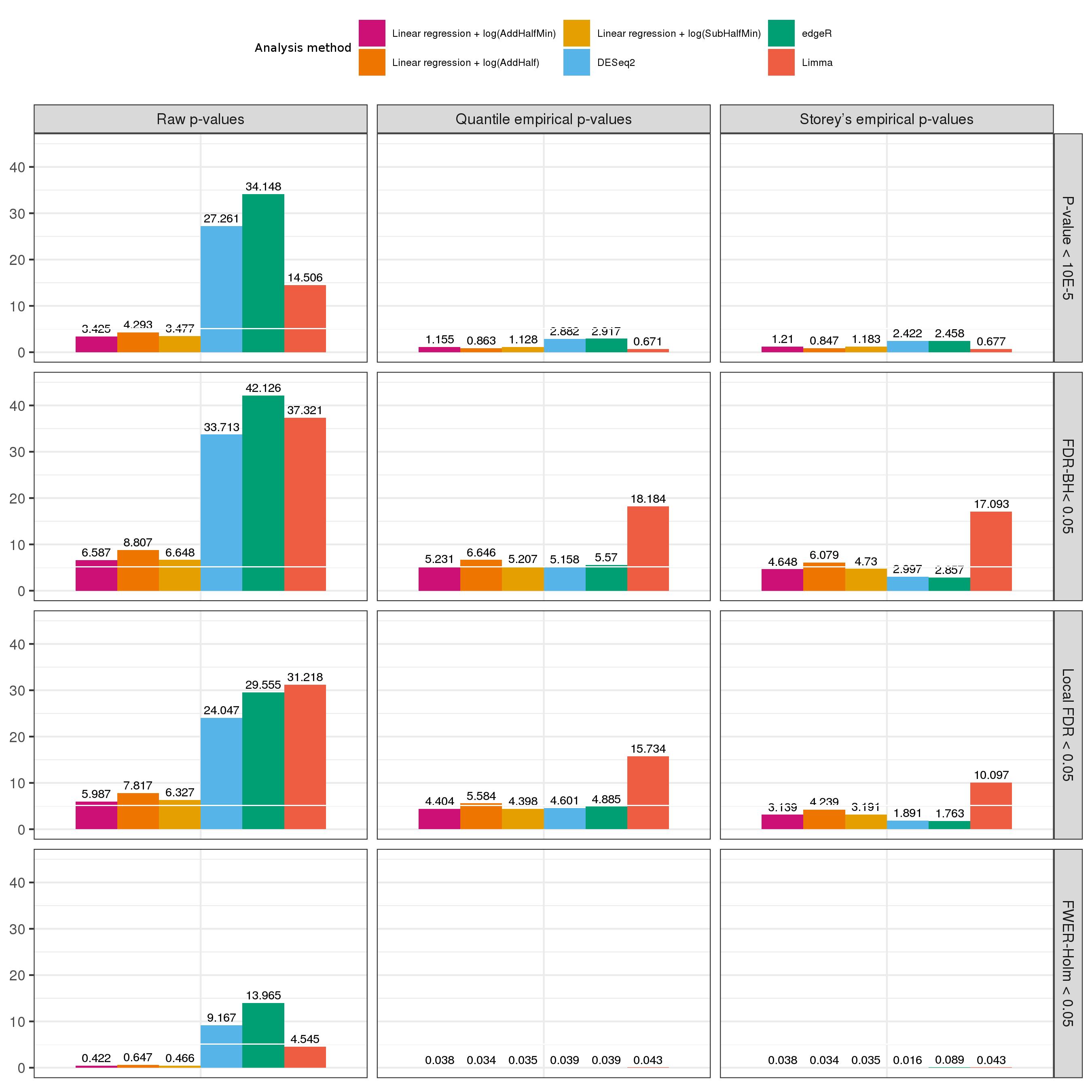

Figure S5: Average number of falsely-detected transcript association with the residual-permuted **MinO2** phenotype, estimated over 100 permutations. We compared approaches based on linear regression, DESeq2, limma, and EdgeR packages; raw p-values, quantile-empirical p-values, and Storey’s empirical p-values; and associations declared as significant according the arbitrary threshold of p-value < ${10}^{-5}$, FDR adjustment using the Benjamini-Hochberg (BH) procedure and using the local FDR procedure implemented in the qvalue R package, and FWER adjustment using Holm procedure.

### Comparison of the effect of normalization methods on false positive detections

We also performed simulation study 1 using, instead of Median normalization, two commonly used normalizations: the TMM normalization implemented in the edgeR package (Figure S6), and the size factor normalization implemented in the DESeq2 package (Figure S7). One can see that the results are very similar to those from Figure S4.

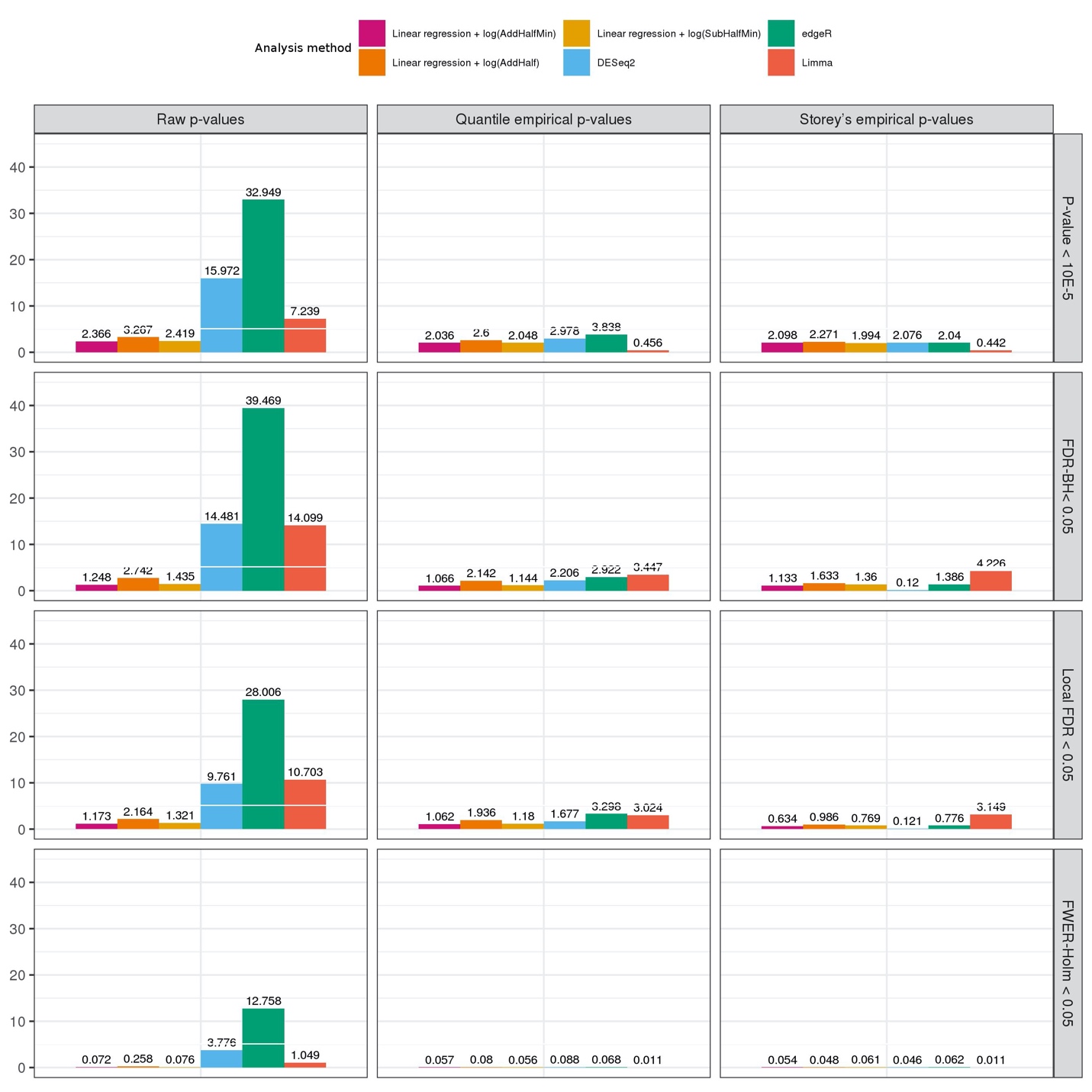

Figure S6: Average number of falsely-detected transcript association with the residual-permuted **AvgO2** phenotype, estimated over 100 permutations. The analysis was applied after **TMM normalization**. We compared approaches based on linear regression, DESeq2, limma, and EdgeR packages; raw p-values, quantile-empirical p-values, and Storey’s empirical p-values; and associations declared as significant according the arbitrary threshold of p-value < ${10}^{-5}$, FDR adjustment using the Benjamini-Hochberg (BH) procedure and using the local FDR procedure implemented in the qvalue R package, and FWER adjustment using Holm procedure.

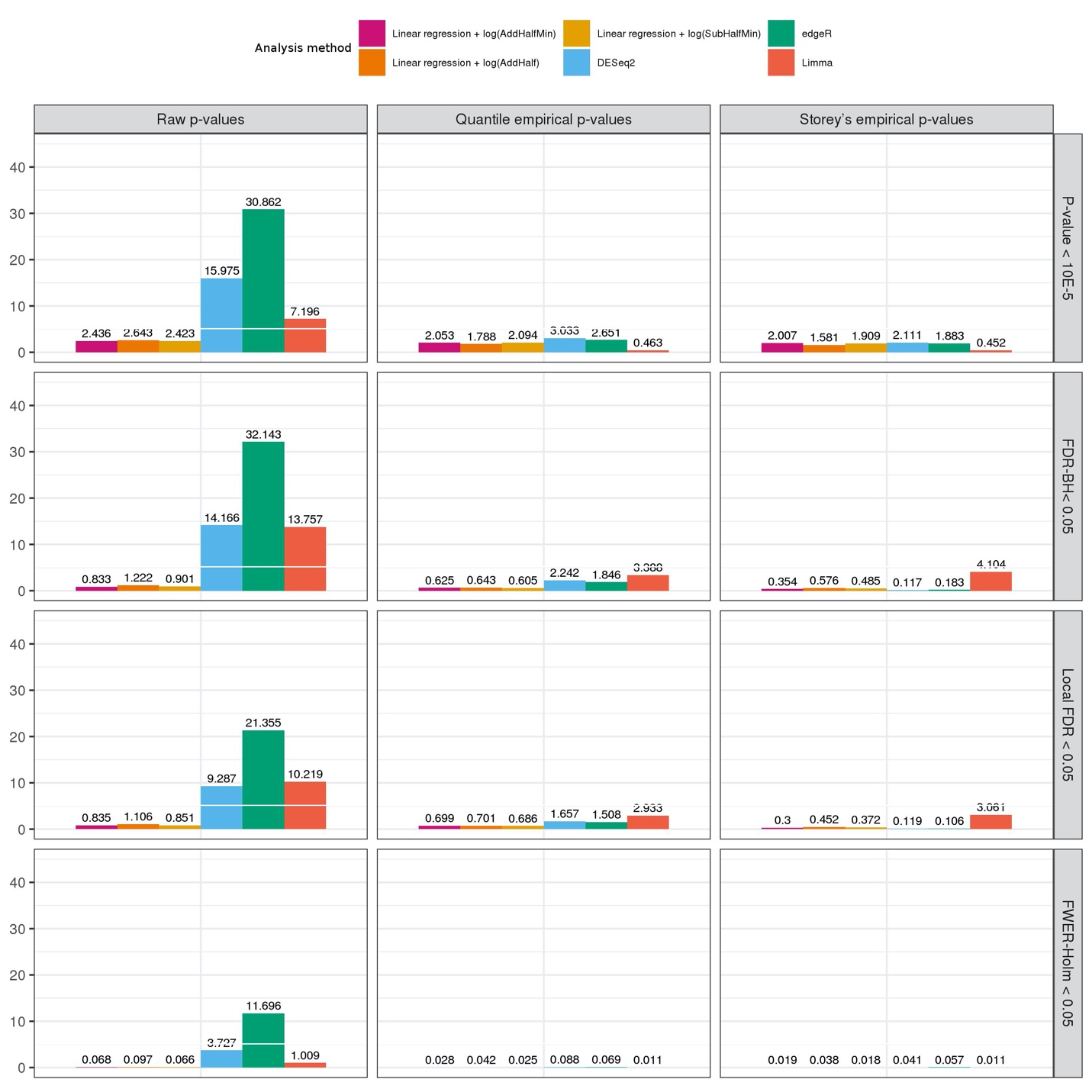

Figure S7: Average number of falsely-detected transcript association with the residual-permuted **AvgO2** phenotype, estimated over 100 permutations. The analysis was applied after **size factor normalization**. We compared approaches based on linear regression, DESeq2, limma, and EdgeR packages; raw p-values, quantile-empirical p-values, and Storey’s empirical p-values; and associations declared as significant according the arbitrary threshold of p-value < ${10}^{-5}$, FDR adjustment using the Benjamini-Hochberg (BH) procedure and using the local FDR procedure implemented in the qvalue R package, and FWER adjustment using Holm procedure.

### Results from simulations study 3: power for detecting an association with a simulated transcript association in a transcriptome-wide association analysis

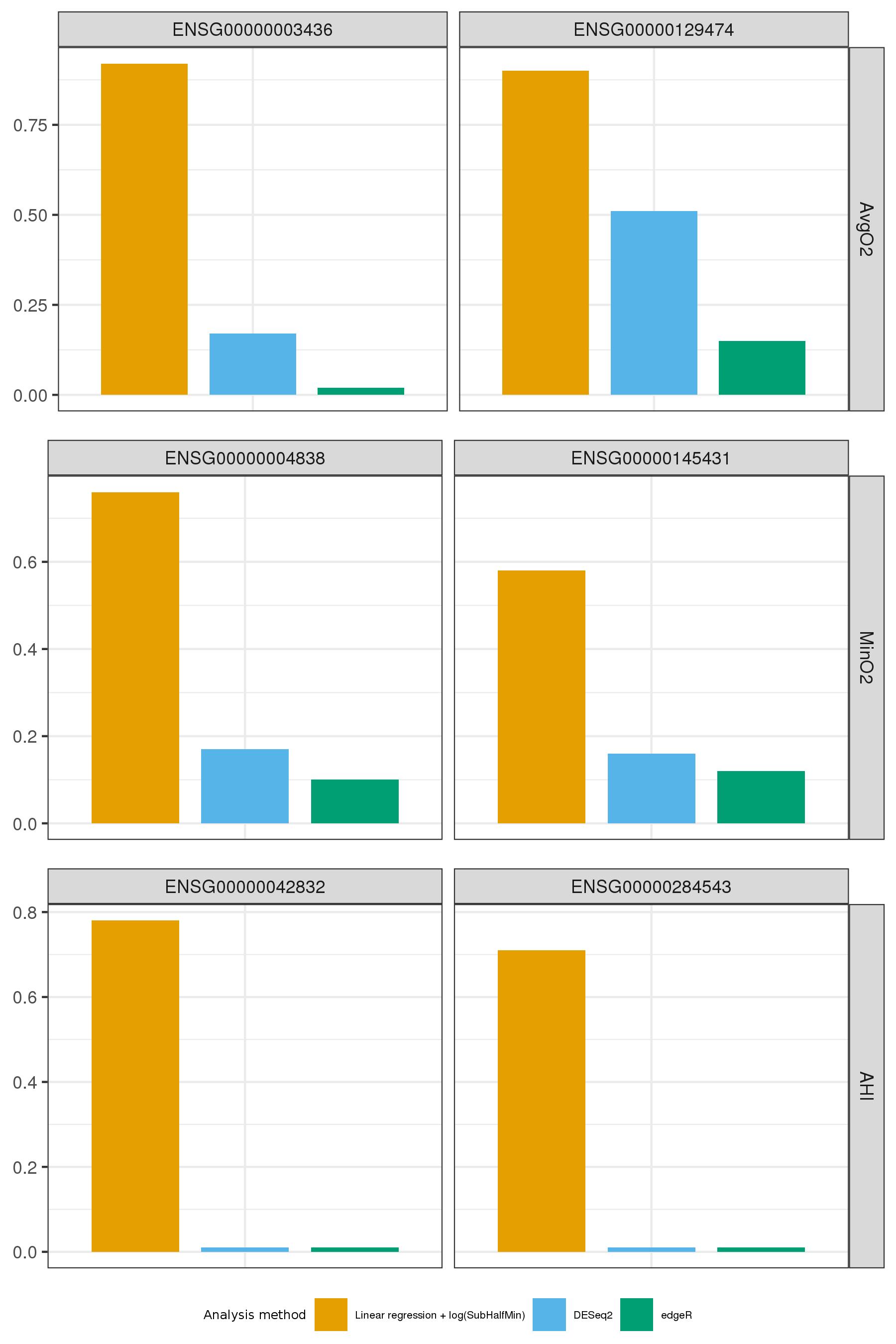

Figure S8: Estimated power for detecting a transcript simulated as associated with the three sleep traits when using **quantile empirical p-values** (compared to Storey empirical p-values in the main manuscript), and association is determined significant if its BH FDR-adjusted p-value is <0.05. We compared logistic regression, DESeq2, and edgeR in transcriptome-wide association analysis for each of the phenotypes.

### Comparison of association discovery using a dichotomized versus a continuous phenotype

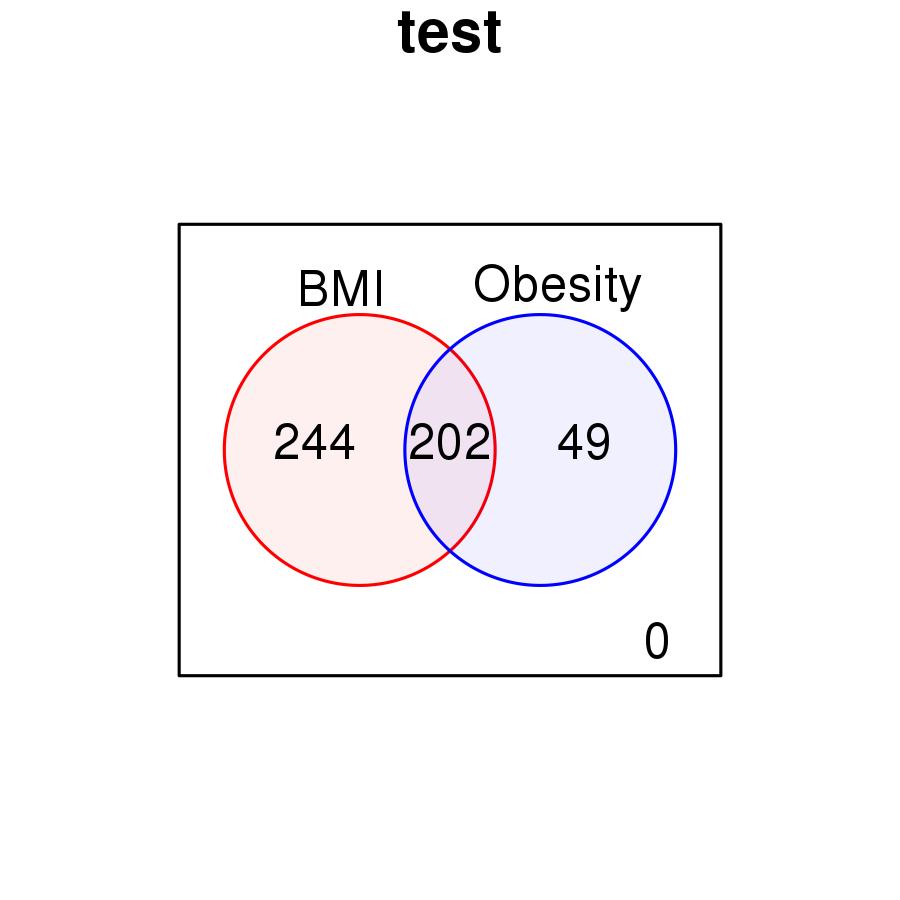

Figure S9: Number of transcripts associated and overlapping between BMI and obesity. Obesity was defined as BMI$\geq30$. We used the same filtering criterion for both phenotypes, requiring 50% non-zero transcript values. For obesity, we also relaxed the filtering and required 20% and 30% non-zero values in other analyses, and the results were similar.

1.	Li B, Dewey CN. 2011. RSEM: accurate transcript quantification from RNA-Seq data with or without a reference genome. *BMC Bioinformatics* 12:323

2.	van der Laan MJ, Hubbard AE. 2006. Quantile-function based null distribution in resampling based multiple testing. *Stat Appl Genet Mol Biol* 5:Article14

3.	Storey J, Bass A, Dabney A, Robinson D. 2019. qvalue: Q-value estimation for false discovery rate control. In *R package version 2.18.0.*
